# Effects of selfing on speciation through the accumulation of Dobzhansky-Muller incompatibilities

**DOI:** 10.1101/2023.03.13.532387

**Authors:** Kuangyi Xu

## Abstract

Phylogenetic analyses suggest that self-compatible lineages have higher speciation rates than self-incompatible lineages. However, the effects of selfing on speciation remain unclear. Although a selfing population can resist gene flow from other populations, selfing may increase gene flow from a focal population to other populations. This study investigates the effects of selfing rates of two populations on the waiting time to speciation through the accumulation of Dobzhansky-Muller incompatibilities (DMI). Generally, a higher mean selfing rate of two populations facilitates speciation when incompatibility-controlling alleles are recessive and are weakly selected, and when gene flow is mainly through pollen dispersal instead of seed dispersal. However, the selfing rate difference between two populations can retard speciation, especially when the selfing rate of immigrants remains unchanged after migration. When the selfing rates of two populations differ, speciation may be fastest when the mean selfing rate is intermediate. Given that selfing rates often vary among conspecific populations in plant species, the results indicate that lineages with higher mean selfing rates may not necessarily have higher rates of speciation through the accumulation of DMI, and also call for an estimation of the dependency of speciation rates on selfing rates.

## Introduction

Evolution from outcrossing to selfing is one of the major evolutionary transitions in plants, which can greatly affect the rate of lineage diversification. Indeed, macroevolutionary analyses show that lineages capable of selfing (i.e., self-compatible) have much higher speciation rates than self-incompatible lineages (Takebayashi and Morrell 2001, Goldberg et al. 2009, Igic and Busch 2013), despite a much higher extinction rate of self-compatible lineages. Nevertheless, from the microevolutionary perspective, speciation can be achieved through various mechanisms (Coyne and Orr 2004), but the effects of selfing on speciation through different mechanisms remain under-investigated.

One of the most important mechanisms of speciation that involves hybrid inviability is the accumulation of Dobzhansky-Muller incompatibilities (DMI). DMI are caused by inappropriate epistatic interactions between alleles at different loci. Compared to other mechanisms (e.g., ecological adaptation), DMI is an intrinsic genetic mechanism and thus depends less on the external environment. Classical models of speciation through the accumulation of DMI often focus on allopatric speciation (e.g., Orr 1995, Turelli and Orr 2000, Yamaguchi and Iwasa 2013), and a substantial amount of later works has investigated the accumulation of DMI under parapatric speciation. Specifically, Gavrilets (2000) showed that migration between populations can greatly increase the expected waiting time to speciation, and later work shows that speciation can be very slow if incompatibilities are severe even at a small difference in genetic distance (Yamaguchi and Iwasa 2017). Also, since the rate of gene flow between populations is critical for parapatric speciation, some studies investigate the maximum migration rate for the maintenance of DMI between two populations (Nosil and Flaxman 2011, Bank et al. 2012, Höllinger and Hermisson 2017, Blanckaert and Hermisson 2018, Blanckaert et al. 2020). Generally, these studies suggest that parapatric speciation becomes much slower or more unlikely as gene flow between populations increases, since it can cause alleles accumulated in one population to get fixed in the other population, which reduces genetic divergence. Correspondingly, one of the explanations for why selfing tends to increase speciation rates is that selfing can reduce gene flow from other populations to the local population (Cutter 2019).

Nevertheless, selfing can have opposing effects on the accumulation of genetic divergence between populations and thus may not necessarily facilitate speciation. First, selfing will increase homozygosity and also reduce the effective population size (Caballero and Hill 1992), and thus influence the fixation probability of incompatibility-controlling alleles. Second, although a selfing population can resist gene flow from other populations, selfing may increase gene flow from a focal population to other populations since selfing immigrants can enjoy a transmission advantage. This can be seen by looking at how immigrant alleles of an immigrant individual over generations. For an outcrossing immigrant, the frequency of immigrant alleles in its offspring will be reduced every generation because half of genes will be from the residents. In contrast, a completely selfing immigrant can transmit all its genes to the offspring, which allows immigrant alleles to persist longer, especially when hybrid offspring have lower fitness due to genetic incompatibilities. As a result, the rates of gene flow will be asymmetric between two populations with different selfing rates. Since there is often great variation of selfing rates among conspecific populations (Whitehead 2018), it is thus unclear whether self-compatible species will always have a higher speciation rate through the accumulation of DMI than self-incompatible species.

Moreover, although self-compatibility tends to facilitate speciation, it is still unclear how the speciation rate changes with the selfing rate, since “self-compatibility” and “selfing rate” are two different concepts. The former is a binary discrete trait while the latter is a continuous trait, and self-compatibility does not necessarily mean a high selfing rate. It may be likely that speciation rates are enhanced only in lineages with intermediate selfing rates, while lineages with too high or too low selfing rates may have lower speciation rates than self-incompatible lineages. Therefore, a further question is that how the selfing rate affects time to speciation through DMI, which remains unclear since previous works often assume random mating populations (e.g., Gavrilets 2000, Yamaguchi and Iwasa 2017, Bank et al. 2012).

In this study, I build population genetic models to investigate the effects of selfing rates of two populations on the expected time to speciation through the accumulation of DMI under both allopatric and parapatric speciation. Briefly, selfing tends to facilitate speciation when incompatibility-controlling alleles are recessive and weakly selected caused by either ecological adaptation or genetic incompatibility, and also when gene flow is mainly through pollen dispersal rather than seed dispersal. However, when the selfing rates of the two populations differ, speciation may be fastest when the mean selfing rate is intermediate. Also, a larger selfing rate difference between two populations can retard speciation, since gene flow from the population with a higher selfing rate to the other population will be high, especially when the selfing rate of immigrants remains unchanged after migration. Since selfing rates of conspecific populations often vary a lot in plant species, especially those with intermediate mean selfing rates, selfing may not necessarily increase the rate of speciation through the accumulation of DMI. The results suggest that an estimation of the dependence of speciation rates on selfing rates may offer a better understanding of the effects of mating systems on lineage diversification.

### Model Framework

I consider two diploid populations 1 and 2 with an equal constant population size *N* and non-overlapping generations. Key symbols used in the model are summarized in Table 1. The selfing rates of population 1 and 2 are denoted by σ_1_ and σ_2_, respectively. To investigate the effects of the variation and the mean selfing rate on the expected waiting time to speciation, the selfing rates of the two populations can be alternatively expressed by the mean selfing rate 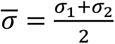, and the selfing rate difference between the two populations 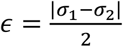. Gene flow between the two populations can be through both pollen and seed dispersal with rates *m*_*p*_ and *m*_*s*_, respectively.

**Table 1.**
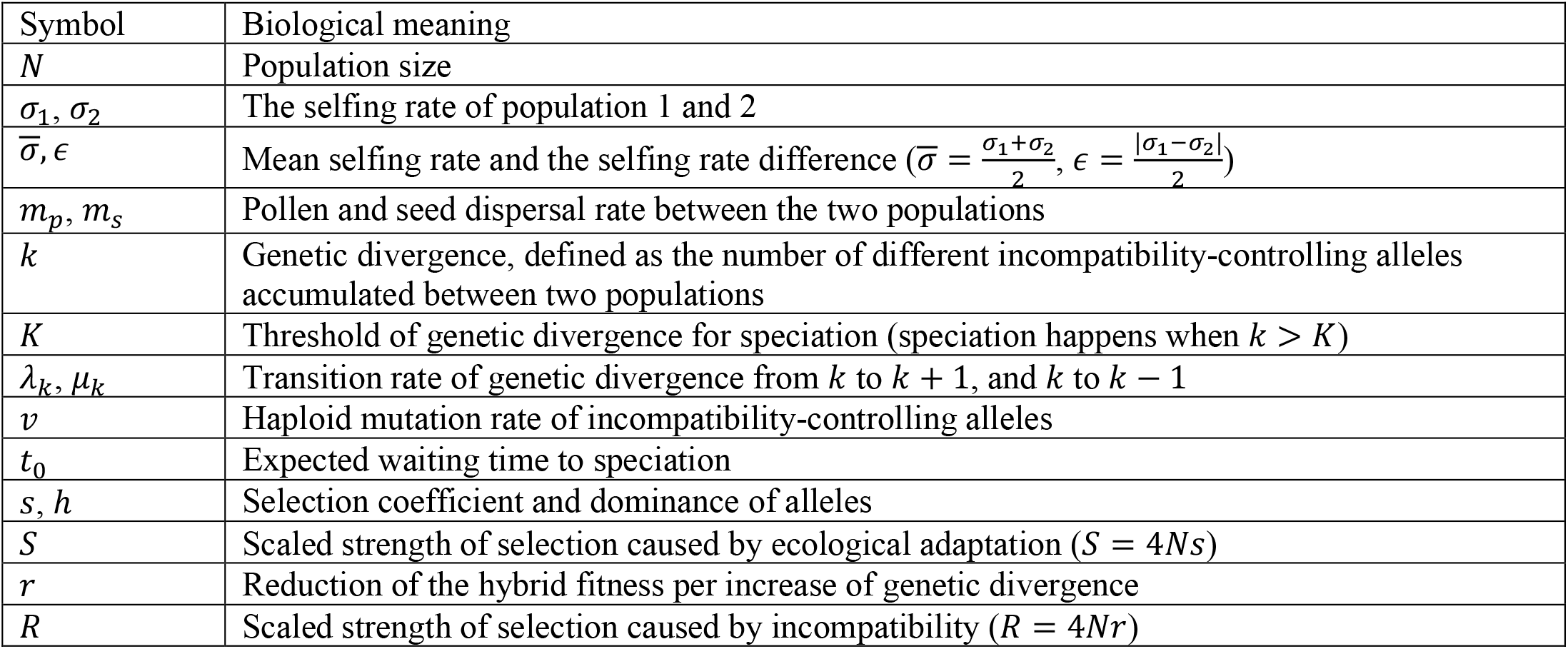
Biological meanings of key symbols.

I consider both allopatric speciation and parapatric speciation. For both types of speciation, I focus on two cases depending on whether alleles are neutral, or are selected for locally ecological adaptation. When alleles are under selection, I assume they are favored in the local population while are selected against in the other population, with the same selection coefficient *s* and dominance *h*, and the loci act multiplicatively on individual fitness in terms of ecological effects.

As will be shown later, the effects of selfing on speciation will depend on how the selfing rate of immigrant individuals changes after dispersal. Therefore, I explicitly distinguish two extreme cases. In the first case, the selfing rate of immigrants will quickly change to that of the local population. This situation may happen if the selfing rate is plastic and is mainly determined by environment (Levin 2012). On the other extreme, I assume the selfing rate of immigrants remains unchanged (Fishman et al. 2002, Goodwillie et al. 2006). In reality, the selfing rate of immigrants may gradually approach that of the local population over time, and we expect the results to be intermediate between the two extreme cases.

I adopt Gavrilets’ model (Gavrilets 2000, Gavrilets et al. 2000, Yamaguchi and Iwasa 2017) for speciation by DMI, which models speciation as a biased random walk with absorption. Instead of directly modelling DMI interactions among loci (Orr 1995, Turelli and Orr 2000), the model focuses on genetic divergence, which is defined as the number of incompatibility-controlling alleles (hereafter referred as alleles for conciseness) that differ between two populations. Genetic divergence increases through the fixation of new mutations in the two populations, but decreases when immigrant alleles from the other population become fixed in the local population. The number of loci is assumed to be infinite so that new mutations will happen at different loci. The haploid mutation rate is *v*, so the overall number of new mutations in a population per generation is 2*Nv*. Also, I assume the expected time for an allele to sweep to fixation is shorter than the expected time for another new mutation to occur, so that interference between alleles is ignored (specifically, if the allele is neutral, the waiting time for a new mutation to arise is 1/*v* and the expected time to fixation is 4*N*_*e*_, so this assumption requires 4*N*_*e*_ *<* 1/*v*). The relative fitness of hybrid caused by genetic incompatibilities, *w*_*h*_(*k*), is assumed to be a non-increasing function of the level of genetic divergence *k*. Speciation is considered to occur when genetic divergence exceeds a threshold number *K*, which means the hybrid is completely inviable (*w*_*h*_(*k*) = 0) when *k* > *K*. During each generation, for *k* ≤ *K*, the probabilities for genetic divergence to increase from *k* to *k* + 1, and decrease from *k* to *k* − 1 are denoted by λ_*k*_ and μ_*k*_, respectively. The haploid mutation rate *v* and the seed and pollen dispersal rates, *m*_*s*_ and *m*_*p*_, are assumed to be small, so that the probability for genetic divergence to increase or decrease by more than 1 is negligibly small.

### Allopatric speciation

#### No selection for local adaptation

Under allopatric speciation, there is no migration between the two populations. When there is no selection for local adaptation, the fixation probability of each mutation is 1/2*N*. Since there are 2*Nv* mutations occurring at each population per generation, the transition rate of genetic divergence from *k* to *k* + 1 is 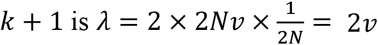. Therefore, the time to speciation is *t*_0_ = (*K* + 1)/2*v* (Gavrilets 2000), which does not depend on the selfing rate.

#### Local adaptation

When alleles are selected for local adaptation, selfing will influence the fixation probability of new mutations. The effective population size of a population decreases with the selfing rate σ as 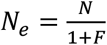 (Caballero and Hill 1992), where 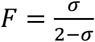 is the inbreeding coefficient, which measures the relative reduction of heterozygosity from that under random mating. The fixation probability of a beneficial mutation in a population

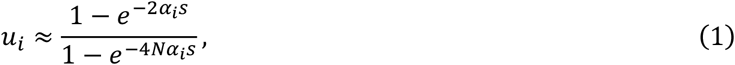

where *α*_*i*_ = (*h* + *F*_*i*_ − *hF*_*i*_)/(1 + *F*_*i*_). The transition rate of genetic divergence from *k* to *k* + 1 is λ_*k*_ = λ = 2*N*(*u*_1_ + *u*_2_), which does not depend on *k*. Given that the population size is large enough (*Nα*_*i*_*s* ≫ 1), the rate of the increase of genetic divergence is approximately 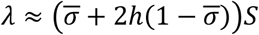, where *S* = 4*Ns*. Therefore, the expected time to speciation is independent of the selfing rate difference between the two populations, *ϵ*. Also, since 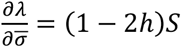, a higher mean selfing rate 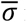 reduces the time to speciation when *h <* 0.5, but increases the time to speciation when *h* > 0.5.

### Parapatric speciation: a threshold hybrid fitness function

In this section, I assume alleles are compatible with each other before genetic divergence *k* reaches the speciation threshold *K*, which means the hybrid fitness function is *w*_*h*_(*k*) = 1 (*k* ≤ *K*). The assumption of a threshold hybrid fitness function is relaxed in the next section.

#### No selection for local adaptation

When alleles are neutral, the transition rate of genetic divergence from *k* to *k* + 1 (*k* ≤ *K*) is the same as that under allopatric speciation, which is λ_*k*_ = 2*v*. The transition rate from *k* to *k* − 1 increases with *k* as μ_*k*_ = *k*μ, where μ is the probability that divergence between the two populations at a locus is lost due to the fixation of immigrant alleles. Therefore, the expected time to speciation is approximately (Gavrilets 2000)

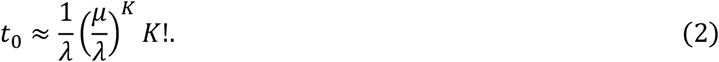

I define the effective migration rate *m*_*e*_ to be the actual proportion of immigrant alleles that enter the local population per generation. Therefore, the total number of migrant alleles between the two populations at each locus per generation is 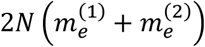, where 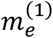 and 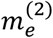 are the effective migration rate from population 1 to population 2 and from population 2 to population 1, respectively. When alleles are neutral, the fixation probability of each allele is 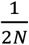, so the probability that divergence at a locus becomes lost is 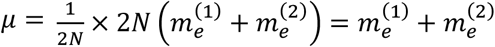. Since 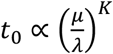 as shown in equation (2), to see the effects of selfing on speciation, it is sufficient to investigate how selfing rates of the two populations affect the effective migration rates.

The overall effective migration rate is the sum of the effective migration rate through pollen and seed dispersal. For gene flow through pollen dispersal, since immigrant alleles enter a local population by fertilizing outcrossed ovules, the effective migration rate through pollen dispersal is 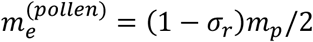, where σ is the selfing rate of the local population. To calculate the effective migration through seed dispersal, I treat immigrants and residents as two categories since their selfing rates differ, and an offspring individual is considered as an immigrant (resident) if the maternal individual is an immigrant (resident). I denote the selfing rates of immigrants and residents by σ_*m*_ and σ_*r*_, respectively, and assume they are constant over time. Over generations, by cross-pollination, immigrant alleles are transmitted to the resident population, and resident alleles also enter the immigrant populations, which dilutes the frequency of immigrant alleles in immigrants. Since the migration rate is assumed to be small, immigrants are rare relative to the residents and almost all outcrossed offspring of immigrants will be fertilized by the residents. Therefore, for an immigrant allele, the number of its offspring alleles in the immigrant population is (1 − σ_*m*_)/2 + σ_*m*_. The two terms capture contributions from outcrossed and selfed offspring, respectively, and the first term is discounted by 1/2 since alleles are only transmitted through female gametes during outcrossing. The number of offspring alleles immigrant allele leaves at generation *t* is [(1 − σ_*m*_)/2 + σ_*m*_]^*t*^. Since the seed dispersal rate is *m*_*s*_, initially there are 2*m*_*s*_ immigrant alleles at a divergent locus. The cumulative number of immigrant alleles over generations is

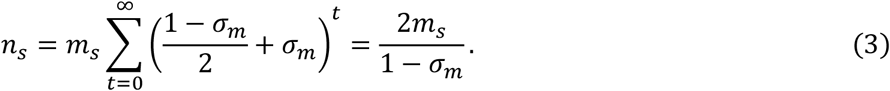

Since the immigrant alleles are transmitted to the resident population through cross-pollination, the effective migration rate caused by seed dispersal, which is the total number of immigrant alleles that enter the resident population, is given by

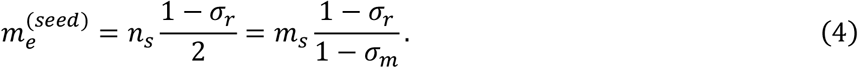

Equation (4) shows that for the special case when 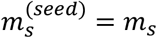. This means when the selfing rates of immigrants and residents are the same, the effective migration rate through seed dispersal is equal to the actual seed dispersal rate, *m*_*s*_. When immigrants have a higher selfing rate than residents (σ_*m*_ > σ_*r*_), the effective migration rate through seed dispersal will be higher than the actual seed dispersal rate (*m*_*se*_ > *m*_*s*_). Particularly, if immigrants are completely selfing (σ_*m*_ = 1), the effective migration rate *m*_*e*_ → ∞ given σ_*r*_ *<* 1. This is because when there is no ecological selection against immigrants, completely selfing immigrants can always persist over generations and disperse pollen to the local population every generation, so that immigrant alleles will finally swamp out the residents through pollen dispersal.

The overall effective migration rate through both pollen and seeds is 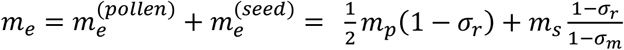. Since how selfing rates of the two populations affect the effective migration rates depends on how the selfing rate of immigrants. Therefore, I consider two extreme cases, depending on whether the selfing rate of immigrants will quickly change to that of the local population, or remains unchanged.

##### The selfing rate of immigrants quickly changes

As previously shown, when the selfing rate of immigrants is the same as the residents, 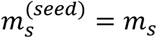.

Therefore, the effective migration rate at the two directions are 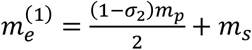, and 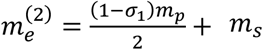, respectively, so the probability for genetic divergence at a locus to get lost is 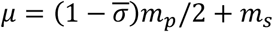. Therefore, a higher mean selfing rate of the two populations, 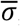, facilitates speciation by reducing gene flow through pollen dispersal. However, the time to speciation does not depend on the selfing rate differences between populations, *ϵ*.

##### The selfing rate of immigrants does not change

When the selfing rate of immigrants remains unchanged, the effective migration rates between the two populations are 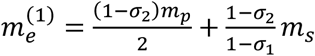, and 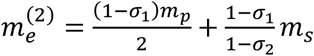. Therefore, the probability that genetic divergence at a locus becomes lost is

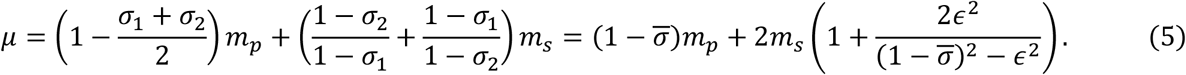

To see the effects of the mean selfing rates 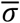 and the selfing rate difference *ϵ* on speciation, since the time to speciation depends on the rate of genetic divergence μ as *t*_0_ ∝ μ^*K*^, it is sufficient to see how the two factors affect μ.

The partial derivatives of μ with respect to 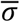 and *ϵ* are

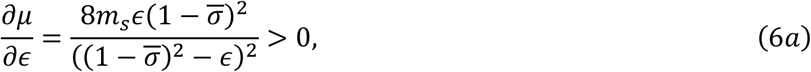

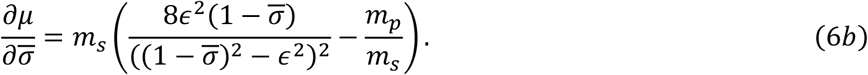

Equation (6a) shows that a greater selfing rate difference between the two populations, *ϵ*, always increases the time to speciation. Equation (6a) shows that the effects of the mean selfing rate 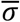 on μ depends on the relative rate of pollen and seed dispersal, and a higher mean selfing rate is likely to reduce μ, and thus the time to speciation, when gene flow is mainly through pollen dispersal (large *m* /*m*). Also, the term 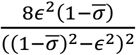 in equation (6b) increases monotonically with a larger 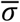, so when 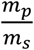 is not too large or small, μ will first decrease and then increase as 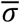 increases. This means that given a fixed selfing rate difference between the two population *ϵ*, there is a certain level of the mean selfing rate 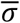 that causes the shortest time to speciation, as illustrated in Fig. 1. This mean selfing rate that leads to the shortest time to speciation decreases as the ratio of pollen dispersal to seed dispersal (*m*_*p*_/*m*_*s*_) becomes lower (Fig. 1(a)), and when the selfing rate difference *ϵ* is larger (Fig. 1(b)).

**Figure 1.**
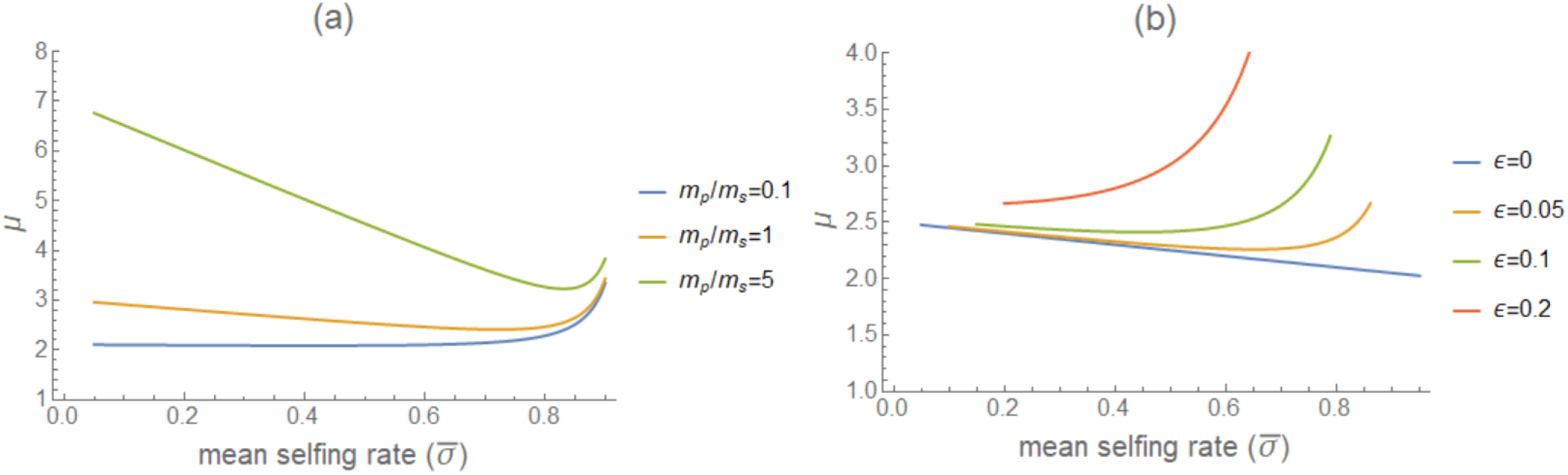
Effects of the mean selfing rate 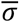 on the probability that genetic divergence becomes lost at a locus μ (note that the time to speciation *t*_0_ ∝ μ^*K*^), when alleles are neutral and the selfing rate of immigrants does not change. In panel (a), the selfing rate difference between populations is *ϵ* = 0.05. In panel (b), the parameter range of the mean selfing rate 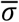 that is biologically feasible is smaller when *ϵ* increases because the selfing rate must range from 0 to 1.

Moreover, as Fig. 1 shows, the mean selfing rate 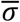 increases, the value of μ, and thus the time to speciation, only changes slightly until 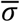 becomes high and μ sharply rises to be high. This is because as 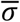 becomes higher, the effective migration rate through seed dispersal from the population with a higher selfing rate (population 2 in Fig. 1) to the other population, 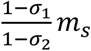, greatly increases.

#### Selection for local adaptation

When alleles are locally favored, the transition rate of genetic divergence from *k* to *k* + 1 is the same as that under allopatric speciation, which is λ_*k*_ = λ = 2*N*(*u*_1_ + *u*_2_), where the fixation probability *u*_1_ and *u*_2_ are given by equation (1). The transition rate of genetic divergence from *k* to *k* − 1, μ_*k*_, is less straightforward, because immigrant alleles are initially completely located on a single haplotype. This linkage disequilibrium among immigrant alleles will gradually decrease through recombination over generations later, which causes the selection intensity at each locus to change over time. However, for tractability, I assume immigrant alleles instantly reach linkage equilibrium at the second generation, which may be a good approximation when the recombination rate is high enough.

To calculate the effective migration rate, when there is only one locus, the number of offspring alleles an immigrant allele leaves to the next generation after selection and reproduction is (1 − *hs*)(1 − σ_*m*_)/2 + (1 − *s*)σ_*m*_. However, when there are multiple loci, the number of offspring alleles at each locus will be affected by selection at other loci. The fitness contributed by a locus at generation *t, w*_*t*_, can be obtained by recursion equations of genotype frequencies. Under linkage equilibrium, the overall fitness of an immigrant individual is the multiplication of *w*_*t*_ across the *k* divergent loci, so the offspring number of an immigrant allele at generation *t* is 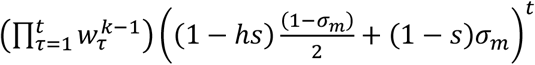. Therefore, the cumulative number of immigrant alleles through seed dispersal is

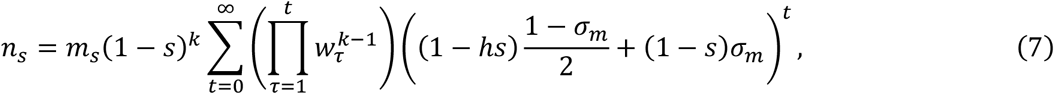

where the term (1 − *s*)^*k*^ accounts for selection at the initial generation when all *k* loci are completely linked. The overall effective migration rate is thus *m*_*e*_ = (*m*_*p*_ + *n*_s_)(1 − σ_*r*_)/2. Note that equation (7) is an overestimation since it only accounts for linkage disequilibrium at the initial generation. However, numeric simulations based on deterministic recursion equations of genotype frequencies [available on Zenodo upon acceptance] suggests that given weak selection, equation (7) is a good approximation (Table S1) for any levels of recombination. Actually, *n*_s_ only slightly changes with the recombination rate between loci. Equation (7) shows that selfing has opposing effects on the cumulative number of immigrant alleles *n*_s_. Selfing prevents immigrant alleles from being lost due to outcrossing (term 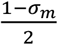 in equation (7)), but also strengthens selection against immigrant alleles by exposing them to homozygotes (term (1 − *s*)σ_*m*_). Although the latter effect is small given weak selection, it can become prominent when there are many divergent loci.

##### The selfing rate of immigrants quickly changes

When the selfing rate of immigrants quickly changes to be the same as the local population, the cumulative number of immigrant alleles from population 1 to population 2, 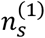, can be calculated by equation (7) by substituting σ_*m*_ with σ_2_ (similarly, σ_*m*_ = σ_1_ for 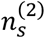). The effective migration rates through both pollen and seed dispersal between populations are 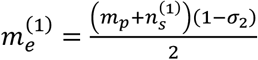, and 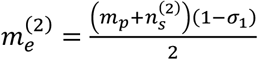. The probability for genetic divergence at a locus to become lost is

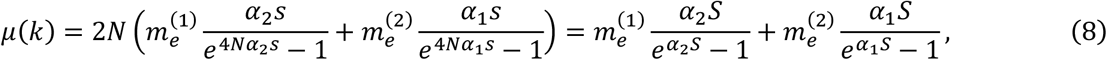

where the term 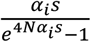 is the fixation probability of an immigrant allele. Unlike the case when alleles are neutral, μ is no longer constant but depends on the level of genetic divergence *k*. Due to the dependency of μ on genetic divergence *k*, the expected waiting time to speciation should be calculated based on a more general expression (Gavrilets 2000) as

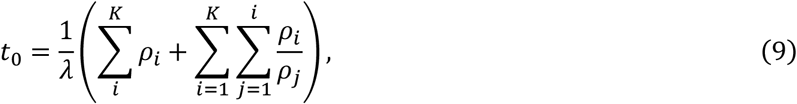

where 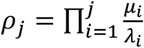

Fig. 2 shows the effects of the mean selfing rate and the dominance coefficient on the time to speciation in the log scale under different levels of selection intensity *S* = 4*Ns*. When selection is weak (*S* = 0.1), a higher mean selfing rate 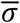 decreases the time to speciation even when alleles are dominant. This is expected since results under weak selection should be similar to the results when alleles are neutral. In contrast, under relatively strong selection (*S* = 10), approximately, a higher 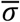 facilitates speciation only when *h* is lower than 0.5.

**Figure 2.**
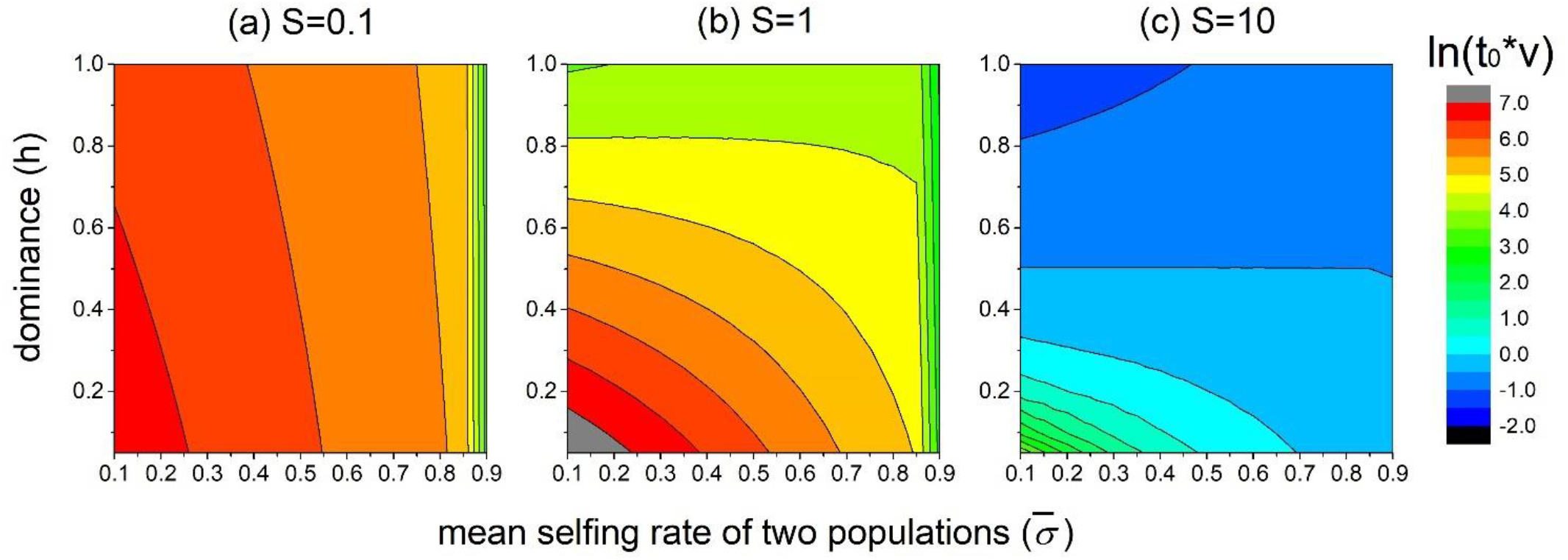
Effects of the mean selfing rate 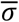 and dominance *h* of alleles on the waiting time to speciation *t*_0_ given a fixed selfing rate difference between populations *ϵ*, when alleles are selected for local adaptation and the selfing rate of immigrants quickly changes. The time to speciation *t*_0_ is scaled by the mutation rate *v* (i.e., *t*_0_*v*) and is shown in the log scale. Parameters are *m*_*p*_ = *m*_*s*_ = *v, ϵ* = 0.1, *K* = 5.

Fig. 3 shows how selfing rates of the two populations σ_1_ and σ_2_ affect the relative time to speciation compared to the situation when both populations are completely outcrossing (σ_1_ = σ_2_ = 0). Generally, a higher selfing rate of either population tends to lower the time to speciation when alleles are recessive (Figs. 3(a), 3(c)), while retards speciation when alleles are dominant (*h* = 0.8, Figs. 3(c), 3(f)). Also, selfing is more likely to facilitate speciation when the divergence threshold for speciation, *K*, is higher, which can be seen by referring to the case when alleles are additive. Specifically, as the mean selfing rate increases, the relative time to speciation only slightly decreases when *K* = 5 unless one of the two populations is highly selfing (Fig. 3(b)), while the decrease of the time to speciation becomes much more prominent when *K* = 20 in Fig. 3(e).

**Figure 3.**
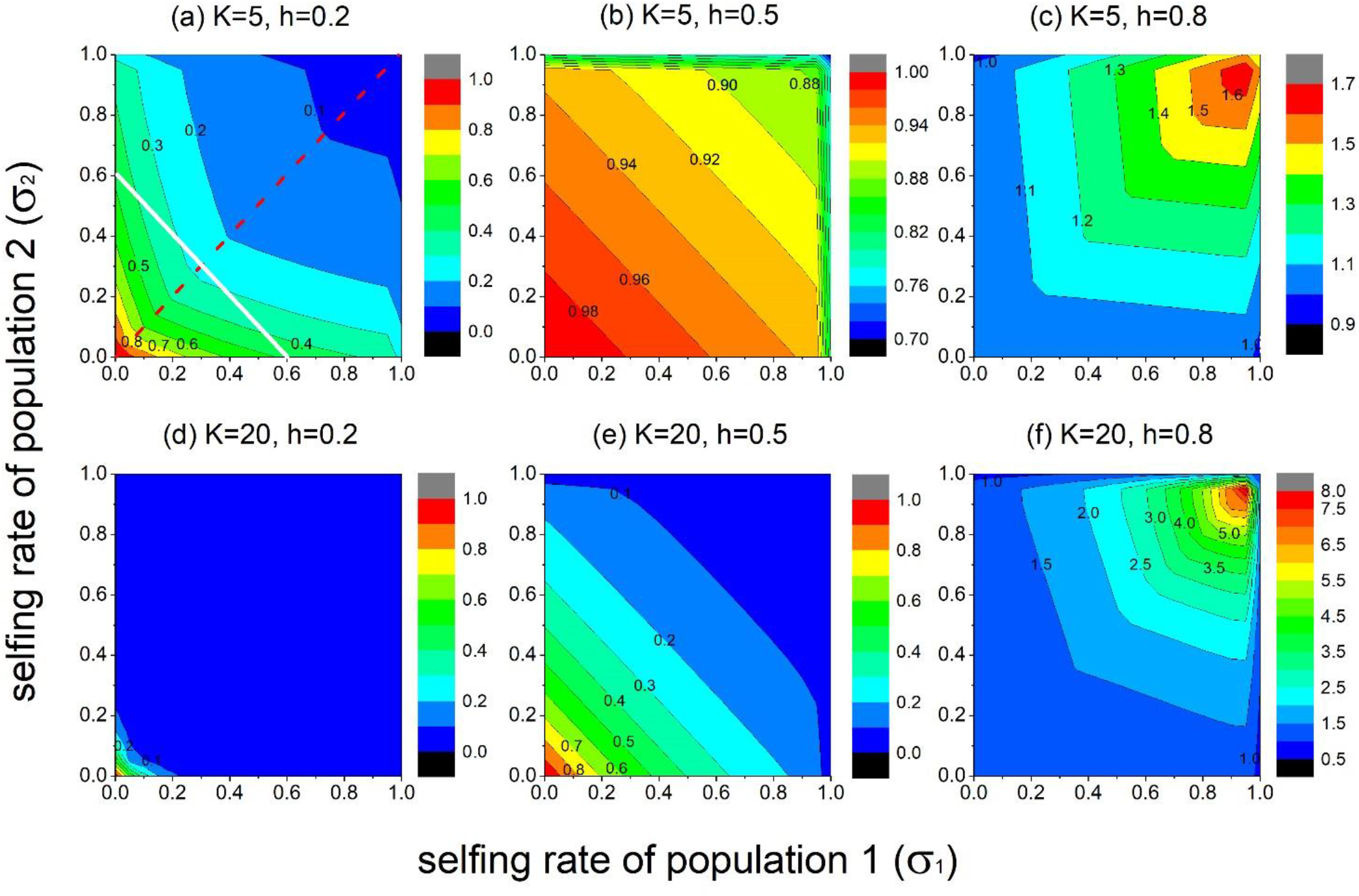
Effects of the selfing rates of the two populations, σ_1_ and σ_2_, on the relative time to speciation compared to that between two completely outcrossing populations (i.e., relative 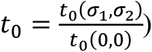, when alleles are selected for local adaptation and the selfing rate of immigrants quickly changes. Panels are symmetric at the diagonal since the two populations are assumed to be identical in other parameters except their selfing rates. Other parameters are *m*_*p*_ = *m*_*s*_ = *v, S* = 5.

The effects of the selfing rate difference between populations, *ϵ*, also depends on dominance of alleles. In Fig. 3, points along a line with slope -1 (e.g., the white line in Fig. 3(a)) have the same mean selfing rate σ, while the selfing rate difference *ϵ* is increases from 0 when points deviate away from the diagonal (the red line in Fig. 3(a)). When alleles are recessive (Fig. 3(a)), a larger *ϵ* tends to increase the time to speciation. However, even when *ϵ* is large, the relative time to speciation is smaller than 1. This means that for two self-compatible populations, even when their selfing rates differ a lot, speciation is still faster than two completely outcrossing populations. When alleles are additive (Figs. 3(b), 3(e)), *ϵ* has nearly no effect on the time to speciation. When alleles are dominant, as Fig. 3(c) and 3(f) illustrate, a larger *ϵ* tends to facilitate speciation, but the relative time to speciation is generally larger than 1, so that speciation between self-compatible populations is slower than completely outcrossing populations.

The effects of selfing on speciation also depends on the relative rate of pollen and seed dispersal, *m*_*p*_/*m*_*s*_. Selfing is more likely to facilitate speciation when gene flow is mainly through pollen dispersal (large *m*_*p*_/*m*_*s*_), since a higher selfing rate of either population always reduces gene flow through pollen dispersal, but not for gene flow through seed dispersal. In fact, under large *m*_*p*_/*m*_*s*_, when alleles are dominant, the time to speciation no longer monotonically increases with a higher mean selfing rate 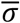, but will first increase and then decrease, which means speciation is slowest when the mean selfing rate is intermediate (see Fig. S1).

##### The selfing rate of immigrants does not change

When the selfing rate of immigrants remains unchanged, the effective migration rates become 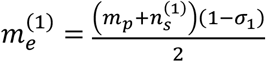, and 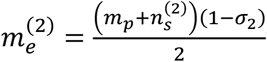. The time to speciation can still be calculated by equations (8) and (9). In contrast to the case when the selfing rate of immigrants quickly changes, when the selfing rate of immigrants remains unchanged, the time to speciation can change dramatically with selfing rates σ_1_ and σ_2_. As Fig. 4 shows, given a fixed selfing rate difference *ϵ*, under weak selection (*S* = 0.1 in Figs. 4(a), *S* = 1 in Fig. 4(b)), the time to speciation tends to be shortest when the mean selfing rate 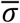 is intermediate. When selection is strong (*S* = 10 in Fig. 4(c)), the time to speciation *t*_0_ increases with a higher mean selfing 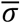 approximately when *h* > 0.5, while decreases with a higher 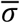 when *h <* 0.5. However, in all panels of Fig. 4, *t*_0_ dramatically increases as 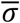 becomes very high, because the effective migration rate becomes very large (see equation (4)).

**Figure 4.**
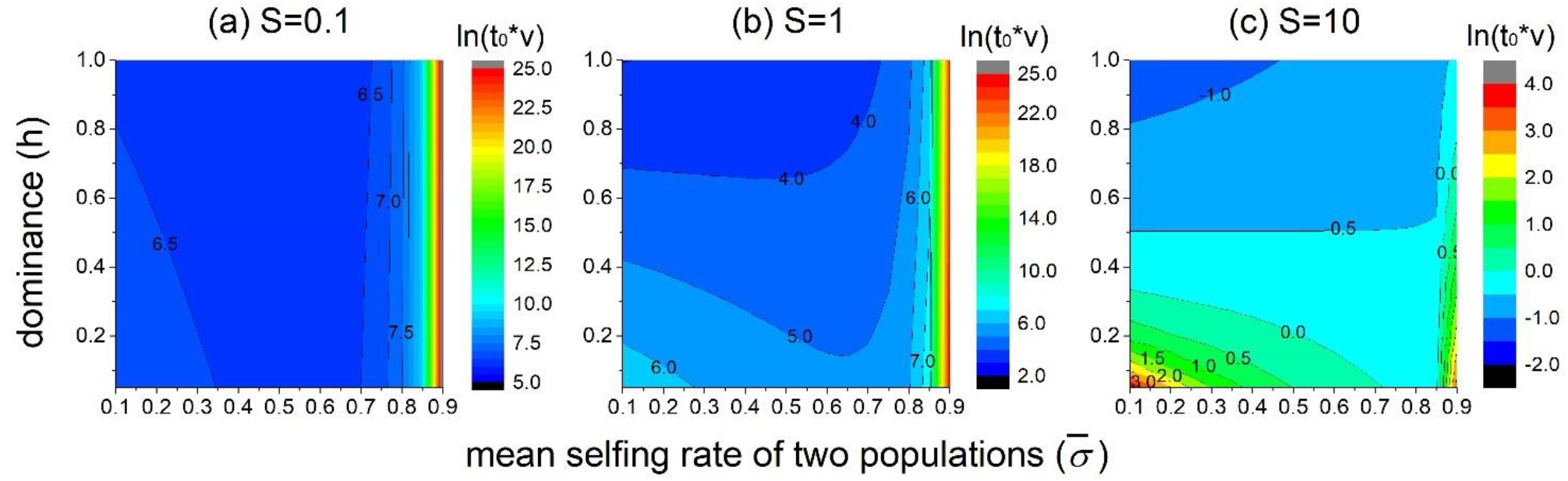
Effects of the mean selfing rate 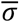 and dominance *h* on the expected time to speciation *t*_0_ given a fixed level of the selfing rate difference between two populations *ϵ* when alleles are selected for local adaptation and the selfing rate of immigrants does not change. Parameters are *m*_*p*_ = *m*_*s*_ = *v, ϵ* = 0.1, *K* = 5.

Fig. 5 illustrates that similar to the results when alleles are neutral, the selfing rate difference *ϵ* tends to increase the time to speciation *t*_0_ for all levels of dominance. As a result, even when alleles are recessive, populations with a high mean selfing rate but a large selfing rate difference may have a longer wating time to speciation than two competely outcrossing populations, which is not found in Fig. 3 where the selfing rate of immigrants quickly changes. When selection is weak, the results are close to the case when alelles are neutral. When selection is intermediately strong, a larger *ϵ* may facilitate speciation is when the speciation threshold *K* is low and when alleles are dominant (*K* = 5, *S* = 5, *h* = 0.8 in Fig. 5(c)), especially when gene flow is mainly through pollen dispersal (*m*_*p*_/*m*_*s*_ large, see Fig. S2). Nevertheless, when selection is strong (*S* = 10, *h* = 0.8, Fig. 5(f)), the time to speciation increases monotonically with a larger *ϵ*. Interestingly, in this case, the time to specication is mainly determined by the lower selfing rate (i.e., min{σ_1_, σ_2_}). For example, in Fig. 5(f), when σ_2_ = 0.4, the time to speciation almost does not change as σ_1_ increases from 0.4. This pattern is partly because when selection is strong and incompatibility-controlling alleles are dominant, genetic divergence is mainly contributed by the accumulation in the population with a lower selfing rate. Also, gene flow from the population with a higher selfing rate to the population with a lower selfing rate no longer influences the time to specation, because most immigrant alleles will be eliminated in the initial several generations after migration.

**Figure 5.**
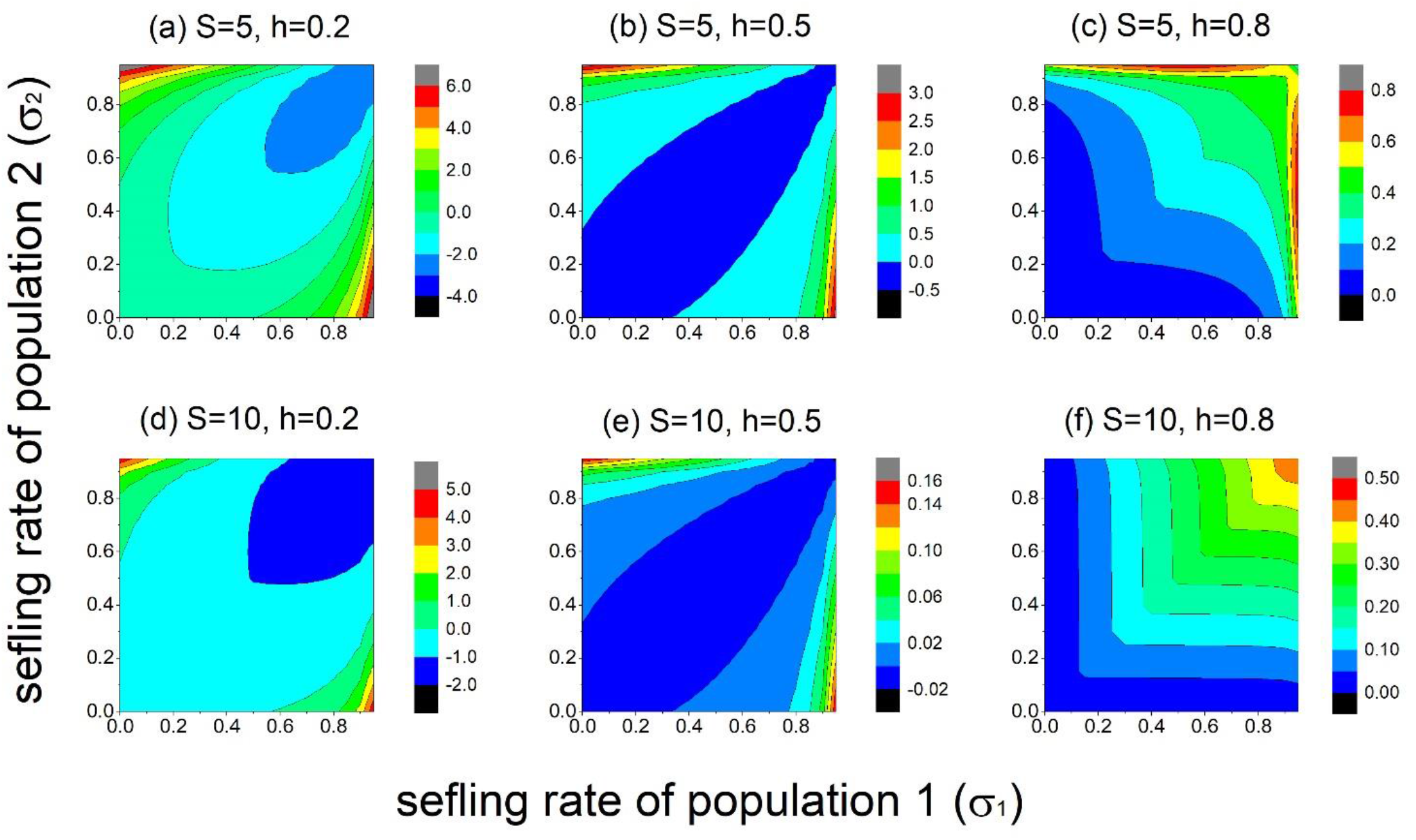
Effects of the selfing rates of the two populations on the waiting time to speciation compared to the between two completely outcrossing populations, when alleles are selected for local adaptation and the selfing rate of immigrants does not change. The relative time to speciation is shown in the log scale. Parameters are *m*_*p*_ = *m*_*s*_ = *v, K* = 5.

### Parapatric speciation: hybrid fitness decreases linearly with genetic divergence

In the previous section, I assume the fitness of hybrids remains constant until genetic divergence exceeds the speciation threshold. However, the fitness of hybrids may become lower as genetic divergence increases. Therefore, here I assume the fitness reduction of hybrid increases linearly with genetic divergence, so that the hybrid fitness function is

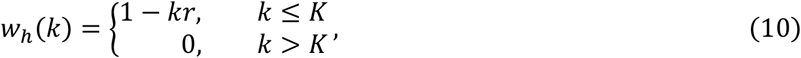

where *r* is the fitness reduction caused by each genetic divergence. Since hybrid fitness depends on the level of genetic divergence, heterozygotes have lower fitness than homozygotes and thus selection on immigrant alleles is frequency-dependent. In the Appendix, I show that the fixation probability of an immigrant allele in a non-completely selfing population is

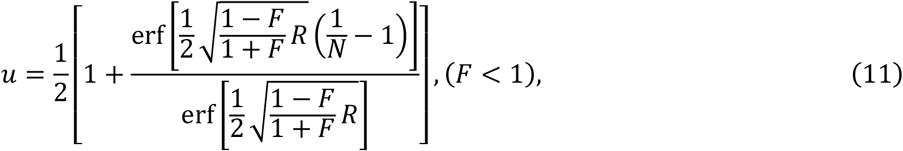

where *R* = 4*Nr* is a scaled parameter that measures the selective strength caused by genetic incompatibilities. The fixation probability *u* is higher when the inbreeding level is higher (i.e., larger *F*), which suggests that a higher mean selfing rate 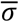 may increase the time to speciation. For a completely selfing population (i.e., *F* = 1), immigrant alleles will always be in homozygotes and thus are neutral, so the fixation probability is *u* = 1/2*N*.

To calculate the cumulative number of immigrant alleles at each locus per generation, note that at the first generation of reproduction, fitness of outcrossed offspring is 1 − *kr* since all immigrant alleles are located on the same haplotype, and fitness of selfed offspring is 1. Therefore, the number of immigrant alleles left is 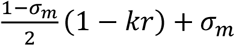. Again, I assume loci instantly reach linkage equilibrium at the second generation, so that the cumulative number of immigrant alleles at a locus is approximately

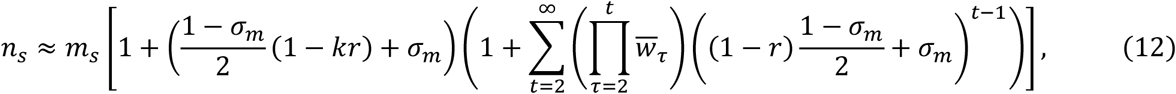

where the mean fitness *w*_*t*_ is obtained by recursion equations of genotype frequencies. Numeric simulation based on deterministic recursion equations of genotype frequencies suggest equation (12) is a good approximation (Table S2). A higher selfing rate of immigrants increases the effective migration rate by allowing immigrants to avoid hybridization with residents. The effective migration rate is again given by *m*_*e*_ = (*m*_*p*_ + *n*_s_)(1 − σ_*r*_)/2, and the time to speciation can be calculated based on equations (8) and (9).

Fig. 6 shows that the effects of selfing on the expected time to speciation mainly depend on the strength of selection caused by genetic incompatibilities, *R*. Generally, a higher mean selfing rate 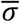 reduces the time to speciation *t*_0_ when selection caused by incompatibilities is weak, as shown in Fig. 6(a) with *R* = 1, and also in Fig. 6(c) given that the selfing rate difference between the two populations is not large. However, a higher 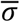 increases *t*_0_ when the fitness reduction caused by incompatibilities is strong, as shown in Figs. 6(b) and 6(d) with *R* = 10. This is because although selfing reduces gene flow, it increases the fixation probability of immigrant alleles (see the text below equation (11)), and the latter effect is more prominent when selection caused by incompatibilities is stronger. The selfing rate difference between populations generally increases the time to speciation, but its effect is slight when the selfing rate of immigrants is the same as the local population (see Fig. 6(b)), while is more prominent when the selfing rate of immigrants remains unchanged, especially when the mean selfing rate is high (see Fig. 6(d)). Similar results are found when the speciation threshold *K* is high (Fig. S3), although the effects of selfing on changing the relative time to speciation become more prominent.

**Figure 6.**
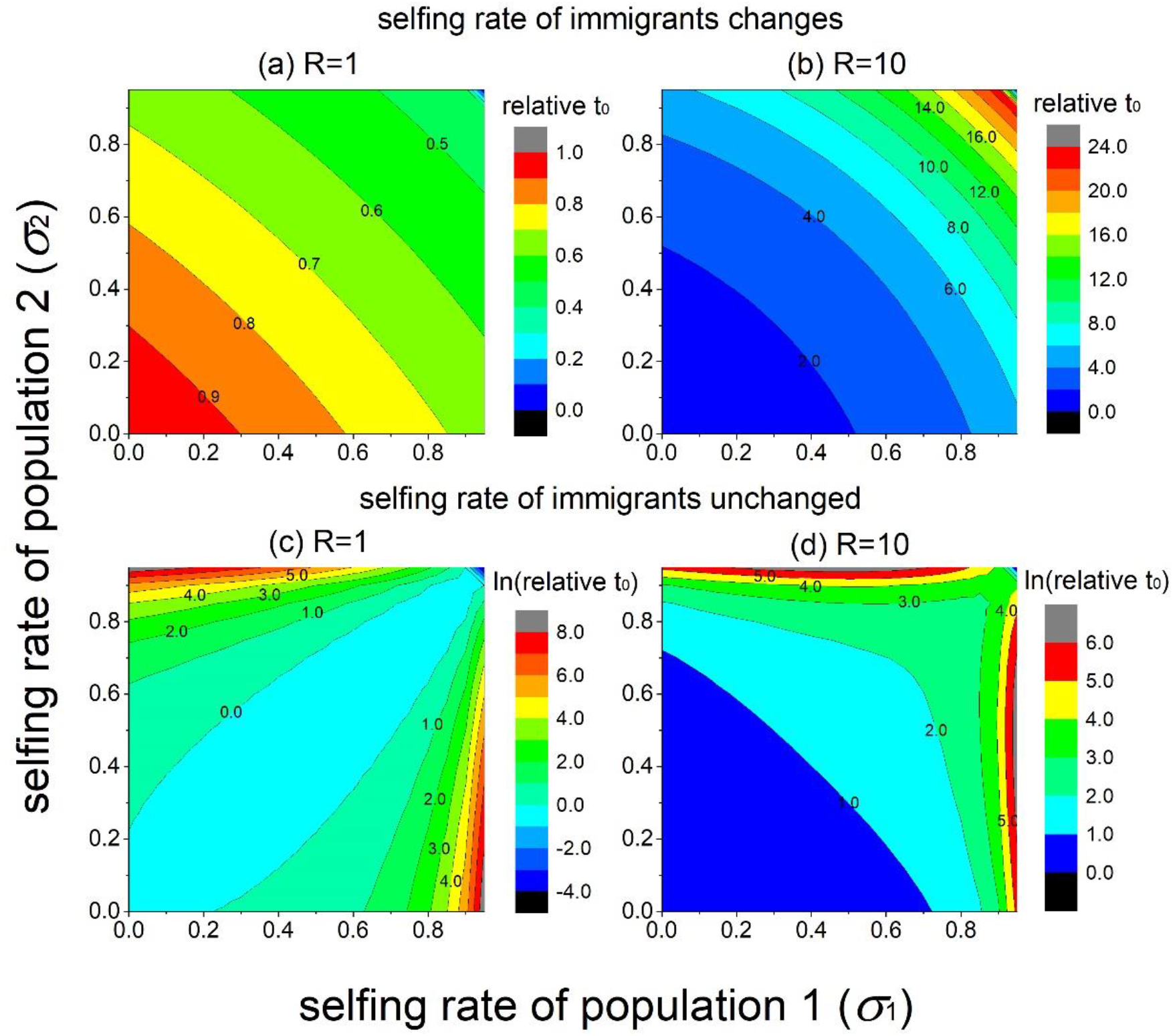
Effects of the selfing rate of the two populations σ_1_ and σ_2_ on the expected time to speciation relative to two completely outcrossing populations when the fitness reduction of hybrids increases with genetic divergence with no local ecological selection. The upper panels show the case when the selfing rate of immigrants quickly changes, and bottom panels show the case when the selfing rate of immigrants does not change. Note that the scale of *t*_0_ differ between upper and bottom panels. Parameters are *m*_*p*_ = *m*_*s*_ = *v, K* = 5.

## Discussion

One important mechanism of speciation is the accumulation of Dobzhansky-Muller incompatibilities (DMI), and previous studies show that the time to speciation through the accumulation of DMI is mainly determined by gene flow between populations (Gavrilets 2000). Although selfing can reduce gene flow from other populations to a focal population, selfing may increase gene flow from a highly selfing population to other populations with a lower selfing rate. This study investigates the effects of selfing on the expected time to speciation through the accumulation of DMI, and the key results are summarized in Table 2. The results identify several key factors that can greatly influence the pattern whether selfing will facilitate speciation or not: 1) the relative contribution of pollen dispersal and seed dispersal, and 2) dominance and the strength of selection on incompatibility-controlling alleles caused by ecological adaptation or genetic incompatibility, and 3) how the selfing rate of immigrant individual changes, and 4) variation of selfing rates among conspecific populations.

**Table 2.**
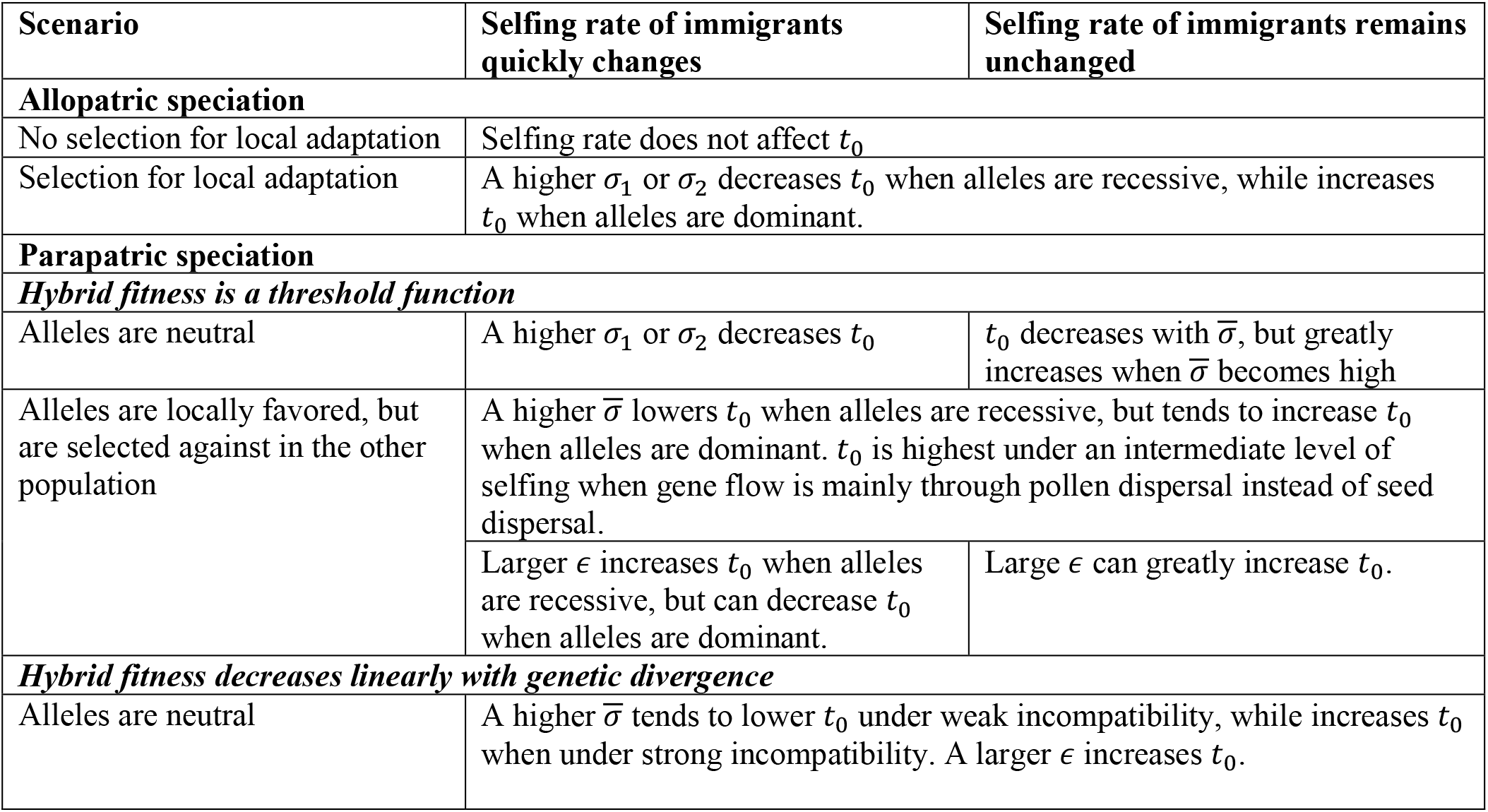
A summary of the effects of selfing rates of two populations σ_1_ and σ_2_ on the expected time to speciation under different scenarios. In the table, 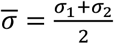 is the mean selfing rate of the two populations, and 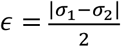 is the selfing rate difference between them. “Alleles” refer to the incompatibility-controlling alleles.

| Scenario | Selfing rate of immigrants quickly changes | Selfing rate of immigrants remains unchanged |
| --- | --- | --- |
| **Allopatric speciation** |  |  |
| No selection for local adaptation | Selfing rate does not affect $t_0$ | |
| Selection for local adaptation | A higher $\sigma_1$ or $\sigma_2$ decreases $t_0$ when alleles are recessive, while increases $t_0$ when alleles are dominant. | |
| **Parapatric speciation** |  |  |
| ***Hybrid fitness is a threshold function*** |  |  |
| Alleles are neutral | A higher $\sigma_1$ or $\sigma_2$ decreases $t_0$ | $t_0$ decreases with $\bar{\sigma}$, but greatly increases when $\bar{\sigma}$ becomes high |
| Alleles are locally favored, but are selected against in the other population | A higher $\bar{\sigma}$ lowers $t_0$ when alleles are recessive, but tends to increase $t_0$ when alleles are dominant. $t_0$ is highest under an intermediate level of selfing when gene flow is mainly through pollen dispersal instead of seed dispersal. | |
| | Larger $\epsilon$ increases $t_0$ when alleles are recessive, but can decrease $t_0$ when alleles are dominant. | Large $\epsilon$ can greatly increase $t_0$. |
| ***Hybrid fitness decreases linearly with genetic divergence*** |  |  |
| Alleles are neutral | A higher $\bar{\sigma}$ tends to lower $t_0$ under weak incompatibility, while increases $t_0$ when under strong incompatibility. A larger $\epsilon$ increases $t_0$. | |

Generally, a higher mean selfing rate tends to facilitate speciation when gene flow is mainly contributed by pollen dispersal instead of seed dispersal, and when incompatibility-controlling alleles are recessive and selection strength of incompatibility-controlling alleles is weak. However, a greater selfing rate difference between populations tends to inhibit speciation, especially when the selfing rate of immigrants does not change after dispersal, mainly due to a high effective migration rate from the highly selfing population to the other population with a lower selfing rate. Indeed, asymmetric introgression from selfing *Mimulus nasutus* populations to outcrossing *Mimulus guttatus* populations has been found in the previous study (Brandvain et al. 2014).

These results may explain why recent studies found no evidence of post-pollination reproductive isolation between self-incompatible and self-compatible populations of *Arabidopsis lyrata* (Gorman et al. 2020, Gorman et al. 2021). Although the current study focuses on speciation through the accumulation of DMI alleles, effects of the asymmetric effective migration rate may also apply to other mechanisms of speciation as long as speciation requires genetic changes that increase divergence between two populations. Although the relative rate of pollen *versus* seed dispersal may vary with species, the migration rate through seeds may be higher than that through pollen if pollinators are limited to the local habitat, which may be likely for long-distance dispersal (Grossenbacher et al. 2017). We may expect pollen dispersal to play a role if pollination is through abiotic vector like wind, or when populations are closer in geography, so that movement of pollinators between populations is possible (Brandvain et al. 2014), and as a result, selfing in these populations should facilitate speciation.

Meta-analysis (Whitehead et al. 2018) suggests that variation of selfing rates among conspecific populations can be large in species with an intermediate mean selfing rate, while is relatively low in predominantly outcrossing or selfing species. In fact, previous models predict that either completely selfing or outcrossing is evolutionarily stable (Lloyd 1979, Lande and Schemske 1985), but populations with mixed mating systems are common (Goodwillie et al. 2005), which may represent a transient stage during evolution from outcrossing to selfing. Based on the fact of great among-population variation in selfing rates, the current results indicate that the speciation rate may become the lowest for species with intermediate selfing rates, while relatively higher in species with low or high selfing rates. Therefore, an estimation of speciation rates based on the mean and among-population variation of selfing rates, instead of self-compatibility, may offer a better understanding of the effects of selfing on speciation.

The current model considered two extreme cases for the selfing rate of immigrants: either quickly changes to be the same as the local population, or remains unchanged. Nevertheless, in reality, the selfing rate of immigrants may gradually approach that of the local population over time, and the actual dynamics should depend on the specific genetic architecture underlying the selfing rate and the pollination environment, but it may be reasonable to expect that the results will be an intermediate between the two extreme cases. Moreover, the model assumes parameters such as the mutation rate and the migration rate to be the same for populations with different selfing rates. However, selfing may also affect the evolution of the dispersal ability and genomic mutation rate, which may influence whether a higher selfing rate will facilitate speciation. The previous model suggests that populations with a higher selfing rate tend to evolve a lower genomic mutation rate (Gervais and Roze 2017).

Also, models show that selfing populations are expected to evolve a lower dispersal rate (Cheptou and Massol 2009, Massol and Cheptou 2011), but self-compatible species may be more successful colonizers by offering reproductive assurance, known as Baker’s law (Baker 1955, see Busch 2011, Pannell et al. 2015 for reviews). The capacity of dispersal may have opposing effects on speciation, since on the one hand, it facilitates establishment of new populations, but on other hand, dispersal between conspecific populations retards speciation. Moreover, selfing may also indirectly affect the rate of gene flow through other factors.

Although rapid speciation associated with the evolution of selfing is found in *Capsella* (Foxe et al. 2009), it should be noted that the study also inferred that the speciation event and the evolution of selfing may happen after colonization events, which results in severe founder effects. Previous models suggest dramatic changes of allele frequencies through genetic drift after a severe founder effect can lead to significant changes in reproductive isolation and thus initiate speciation (Templeton 1980, Templeton 2008, Uyeda et al. 2009), supported by later experiments (Matute 2013). Therefore, selfing may not be the causal effect for rapid speciation, but may be merely a byproduct selected for due to pollen limitation. From this perspective, the higher speciation rate of self-compatible lineages based on the phylogenetic analysis (Goldberg et al. 2010) may be partly due to the fact that self-compatible species is more likely to be successful colonizers, as Baker’s law states. In other words, instead of directly facilitating speciation by reducing gene flow, self-compatibility may be more likely to indirectly facilitate speciation through more frequent successful colonization, which triggers speciation. Given that colonization may play an important role in speciation, it may be interesting for future studies to see how selfing interplays with colonization in affecting speciation.

It should be noted the current study focuses only on speciation through accumulation of DMI, while how selfing overall influences the speciation rate should be understood by studying its effects on speciation through other mechanisms, as well as the relative contribution of these mechanisms to speciation. Therefore, although the current results suggest that selfing may not necessarily promotes speciation through the accumulation of DMI, it is not contradictory to the pattern of higher speciation rates in self-compatible lineages than in self-incompatible lineages (Goldberg et al. 2010). Instead, the results imply that selfing may facilitate speciation mainly through other mechanisms. For example, selfing may be important in facilitating polyploid speciation since selfing helps polyploids establish by avoiding backcross (Baack 2005).

## Appendix

Here I derive the fixation probability of an immigrant allele at a divergent locus when the hybrid fitness is 1 − *r*. Denote the frequency of the immigrant allele as *p*, after one generation, the change of its frequency is

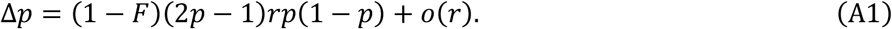

The selection coefficient (1 − *F*)(2*p* − 1)*r* is negative when *p <* 0.5, and is positive when *p* > 0.5. Selfing reduces the magnitude of selection coefficient by increasing inbreeding coefficient *F*, making the allele less unfavored when *p <* 0.5, but also less favored when *p* > 0.5. The fixation probability of an immigrant allele can be calculated by the general expression (Kimura 1962)

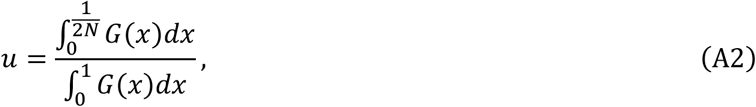

where

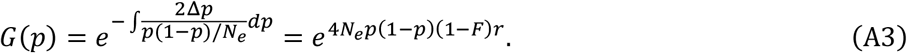

Therefore, the fixation probability of an immigrant allele in a non-completely selfing population is

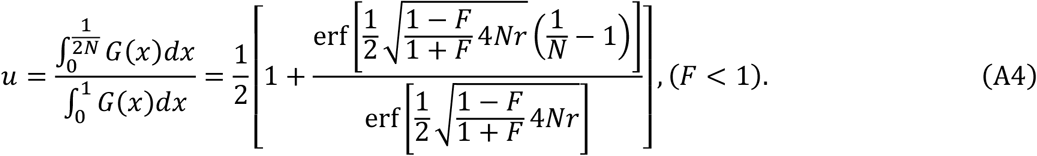

It can be proved that 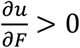, so that the fixation probability *u* increases with inbreeding coefficient *F* (thus a higher selfing rate).

## Supplementary Table and Figures

**Table S1:** Comparison of the cumulative number of immigrant alleles *n*_s_ between results from deterministic recursion equations of genotype frequencies *versus* the approximation by equation (7) in the main text. Loci are assumed to be identical, and “rec” is the recombination rate between two adjacent loci. Rows with “rec” are results from numeric simulations based on deterministic recursion equations. The genetic divergence is *k* = 3. Note that results from equation (7) are scaled by the seed dispersal rate *m*_*s*_ (i.e., *n*_s_/*m*_*s*_).

| Selfing rate | 0 | 0.2 | 0.4 | 0.6 | 0.8 | 1.0 |
| --- | --- | --- | --- | --- | --- | --- |
| **$s = 0.01, h = 0.1$** |  |  |  |  |  |  |
| rec=0 | 1.936 | 2.397 | 3.139 | 4.526 | 8.019 | 32.669 |
| rec=0.01 | 1.936 | 2.398 | 3.140 | 4.529 | 8.035 | 32.669 |
| rec=0.1 | 1.936 | 2.399 | 3.145 | 4.547 | 8.136 | 32.669 |
| rec=0.5 | 1.936 | 2.400 | 3.152 | 4.574 | 8.270 | 32.669 |
| Equation (7) | 1.936 | 2.400 | 3.153 | 4.587 | 8.382 | 32.669 |
| **$s = 0.01, h = 0.9$** |  |  |  |  |  |  |
| rec=0 | 1.898 | 2.343 | 3.057 | 4.391 | 7.766 | 32.669 |
| rec=0.01 | 1.898 | 2.344 | 3.059 | 4.395 | 7.784 | 32.669 |
| rec=0.1 | 1.900 | 2.347 | 3.067 | 4.421 | 7.891 | 32.669 |
| rec=0.5 | 1.901 | 2.351 | 3.082 | 4.458 | 8.038 | 32.669 |
| Equation (7) | 1.901 | 2.351 | 3.082 | 4.472 | 8.145 | 32.669 |

**Table S2:** Comparison of the cumulative number of immigrant alleles between results from exact recursion equations and approximation by equation (12). The strength of incompatibility is *r* = 0.01, and genetic divergence is *k* = 5. Note that results of equation (12) are scaled by *m*_*s*_ (i.e., *n*_s_/*m*_*s*_).

| Selfing rate | 0 | 0.2 | 0.4 | 0.6 | 0.8 |
| --- | --- | --- | --- | --- | --- |
| rec=0 | 1.920 | 2.238 | 3.151 | 4.682 | 9.258 |
| rec=0.01 | 1.936 | 2.239 | 3.168 | 4.717 | 9.348 |
| rec=0.5 | 1.931 | 2.401 | 3.183 | 4.743 | 9.409 |
| Equation (12) | 1.941 | 2.427 | 3.232 | 4.829 | 9.596 |

**Figure S1.**
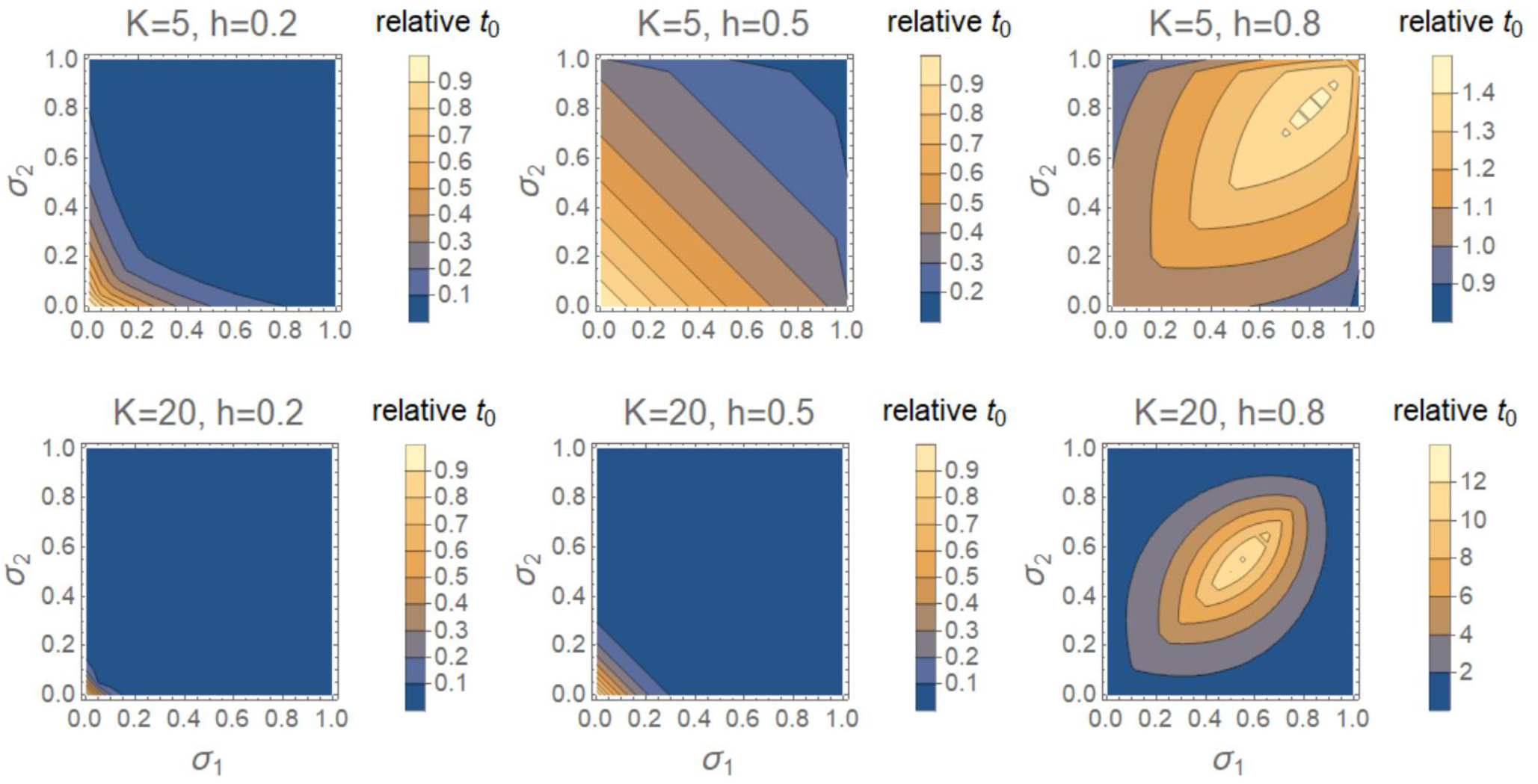
Effects of the selfing rates of the two populations, σ_1_ and σ_2_ on the relative time to speciation compared to that between two completely outcrossing populations (i.e., 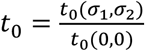 when alleles are locally favored and the selfing rate of immigrants quickly changes. The figure shows the case when the gene flow is mainly through pollen dispersal with parameters being *m*_*p*_ = 10*m*_*s*_ = 10*v, S* = 5. Compared to the case when the gene flow is through both pollen and seed dispersal in Fig. 3 (*m*_*p*_ = *m*_*s*_),

**Figure S2.**
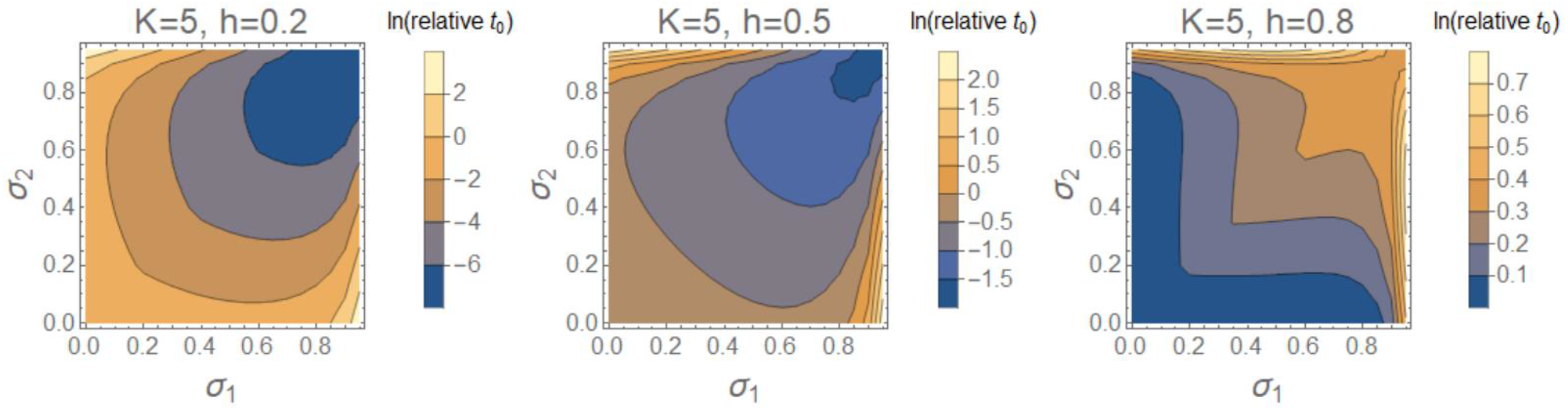
Effects of the selfing rates of the two populations, σ_1_ and σ_2_, on the relative time to speciation compared to that between two completely outcrossing populations when alleles are locally favored and the selfing rate of immigrants does not change. The relative time to speciation is shown in the log scale. Parameters are *S* = 5, *m*_*p*_ = 10*m*_*s*_ = 10*v*.

**Figure S3.**
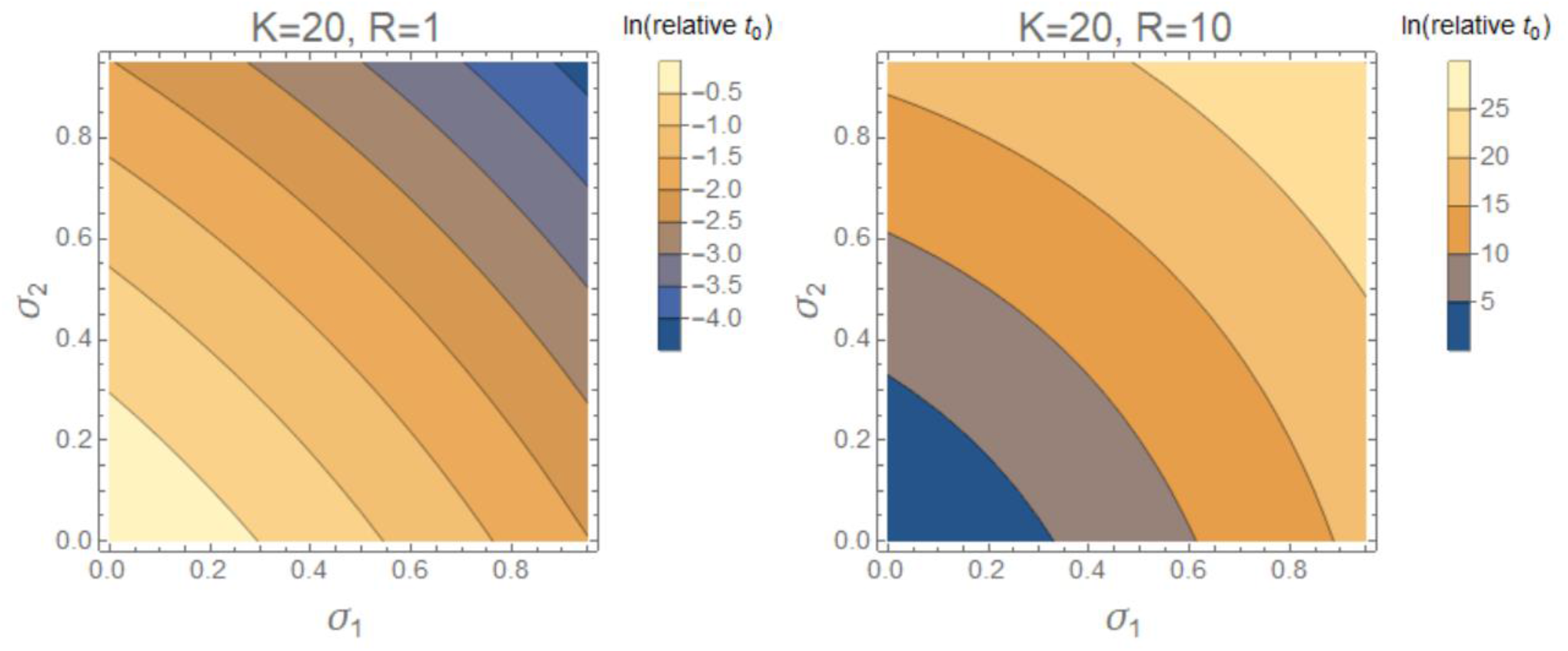
Effects of σ_1_ and σ_2_ on the expected time to speciation relative to that in self-incompatible species when fitness reduction due to reproductive incompatibility increases with genetic divergence under a high speciation threshold *K*.

